# A stable *Netrin-1* fluorescent reporter chicken reveals cell-specific molecular signatures during optic fissure closure

**DOI:** 10.1101/2024.10.01.616014

**Authors:** Brian Ho Ching Chan, Holly Hardy, Teresa Requena, Amy Findlay, Jason Ioannidis, Dominique Meunier, Maria Toms, Mariya Moosajee, Anna Raper, Mike McGrew, Joe Rainger

## Abstract

*NTN1* is expressed in a wide range of developmental tissues and is essential for normal development. Here we describe the generation of a Netrin-1 reporter chicken line (*NTN1-T2A-eGFP*) by targeting green fluorescent protein into the *NTN1* locus using CRISPR/Cas9 methodology. Our strategy gave 100% transmission of heterozygous (*NTN1*^*T2A-eGFP*/+^) embryos in which GFP localisation faithfully replicated endogenous *NTN1* expression in the optic fissure and neural tube floorplate. Furthermore, all *NTN1*^*T2A-eGFP*/+^ embryos and hatched birds appeared phenotypically normal. We applied this resource to a pertinent developmental context - coloboma is a structural eye malformation characterised by failure of epithelial fusion during optic fissure closure (OFC) and *NTN1* is specifically expressed in fusion pioneer cells at the edges of the optic fissure. We therefore optimised the isolation of GFP expressing cells from embryonic *NTN1*^*T2A-eGFP*/+^ eyes using spectral fluorescence cell-sorting and applied transcriptomic profiling of pioneer cells, which revealed multiple new OFC markers and novel pathways for developmental tissue fusion and coloboma. This work provides a novel fluorescent *NTN1* chicken reporter line with broad experimental utility and is the first to directly molecularly characterise pioneer cells during OFC.

## INTRODUCTION

Netrin-1 belongs to a family of secreted extracellular proteins with a diverse range of roles throughout the developing embryo (Chaturvedi and Murray, 2021; Cirulli and Yebra, 2007; Lai Wing Sun et al., 2011). Homozygous loss of functional Netrin-1 during embryonic development causes axon guidance defects (Bin et al., 2015; Serafini et al., 1996; Yung et al., 2015) in addition to tissue fusion-related abnormalities in the eye, inner-ear, and palate (Hardy et al., 2019; Richardson et al., 2019; Salminen et al., 2000a; Yung et al., 2015). Whereas loss of Netrin-1 is lethal in mice, to date only heterozygous missense mutations in *NTN1* have been identified in humans and are associated with congenital mirror movements, a mild disorder of the CNS (Méneret et al., 2017). No phenotypic abnormalities have been reported in heterozygous *Ntn1* targeted mice (Bin et al., 2015; Serafini et al., 1996; Yung et al., 2015).

Netrin-1 has been well-studied for its canonical roles in guidance of commissural and peripheral motor axons and growth-cone dynamics, with attraction or repulsion of axonal projections dependent on the co-expression of specific receptors (reviewed in Lai Wing Sun et al., 2011; Larrieu-Lahargue et al., 2012). Cell-autonomous receptor expression is also associated with the regulation of apoptosis by *NTN1* expressing tissues (Fitamant et al., 2008; Newquist et al., 2013; Nishitani et al., 2017), and Netrins have been associated with tumour metastases, angiogenesis and branching morphogenesis (Chaturvedi and Murray, 2021; Dudgeon et al., 2023; Kye et al., 2004; Lai Wing Sun et al., 2011; Liu et al., 2004). Thus, accurate molecular profiling of *NTN1* expressing and responsive cells would permit in depth analyses for a range of developmental and broader biological processes.

Fusion of opposing sheets of cells during development is critical across all vertebrates, and failure of tissue fusion underlies a range of human developmental disorders, including spina bifida, cleft palate, septal heart defects and hypospadias (Ray and Niswander, 2012). In the developing vertebrate eye, the optic fissure (OF, also referred to as the *choroid fissure*) is a naturally-arising gap extending along the entire proximal-distal length of the early ventral eye cup (Chow and Lang, 2001; Fuhrmann, 2010). Retinal epithelia at the edges of this fissure must completely fuse together to complete the spherical 3D circumferential eye structure and this is required for normal eye development. Incomplete closure of the OF causes ocular coloboma, manifested clinically as persistent open regions of the retina, optic nerve and/or iris, which may severely affect vision (Chang et al., 2006; Gregory-Evans et al., 2004; Patel and Sowden, 2017). OFC involves a range of cell types from distinct origins that contribute either directly or indirectly to the fusion process (Chan et al., 2021; Gestri et al., 2018; Patel and Sowden, 2017), but the recent identification of pioneer cells at the epithelial edges of the OF margins (Eckert et al., 2020; Hardy et al., 2019) offers an opportunity to characterise genes and pathways that specifically regulate fusion. *NTN1* expression was recently shown to be spatially restricted to pioneer cells before and during fusion but was immediately downregulated in the nascently-fused fissure (Hardy et al., 2019; Richardson et al., 2019). Thus, *NTN1* is a pioneer cell marker with specific spatial-temporal expression during fusion phases of OFC that is predicted to tolerate haploinsufficiency during development. We previously characterised OFC in the developing chicken embryo and showed that the size, developmental timing, and ease of segmental dissections within the chicken OFM, were major advantages over other existing models. Coupled to recent advances that have enabled the simplified and expedient generation of gene-edited germline chicken lines for developmental biology (Ballantyne et al., 2021; Woodcock et al., 2017), we set out to make a reporter chicken line by targeting the *NTN1* locus with green fluorescent protein. Our aim was to apply this line to enable the selective isolation and molecular characterisation of pioneer cells to identify new genes and pathways that are active during fusion in OFC and use this approach as an exemplar for this tractable developmental resource.

## RESULTS AND DISCUSSION

### Recombining T2A-eGFP into the chicken *Netrin-1 locus*

Endogenous *NTN1* is expressed in the optic fissure during chicken embryonic development, as well as the floorplate of the neural tube (**Figure 1a**). To develop a *NTN1* fluorescent reporter chicken line, we applied a CRISPR/Cas9 knock-in strategy to deliver a T2A self-cleaving peptide and enhanced GFP (*NTN1-T2A-eGFP*) reporter gene in-frame into the endogenous *NTN1* locus to create a truncated non-functional *NTN1* allele and a cleaved cytoplasmic eGFP from the same locus (**Figure 1b,c**). We first identified sgRNA sequences in *NTN1* exon 1, then cloned these into a CRISPR vector with combined *Cas9* expression and puromycin resistance. We separately generated a repair template construct with the T2A-eGFP open reading frame flanked at both 5’- and 3’-ends with 100 bp flanking homology sequences to the *NTN1* exon 1 cut site. These sequences were further flanked by recognition sequences for the sgRNA to enable high efficiency HDR into the *NTN1* locus (Zhang et al., 2017). We used these combined vectors in co-transfections of male primordial germ cells (PGCs), followed by puromycin selection and FACs to generate single-cell derived clonal cultures for expansion (**Supplemental figure S1**). We identified unambiguous knock-in events in 15 independent PGC lines (15 of 39 analysed clones) by allele-specific PCRs followed by Sanger sequencing; 11 of these were biallelic and 4 were monoallelic (**Figure 1d**; **Supplemental figure S1**). We determined that 100% of all HDR events were in-frame and had no indels or other changes (**Figure 1e**). We injected 12 embryonic day 2.5 iCaspase9 surrogate host chicken embryos with PGCs derived from one single biallelic (homozygous) *NTN1*^*T2A-eGFP/ T2A-eGFP*^ clonal line, according to a previously reported protocol (Ballantyne et al., 2021). Of these injected embryos, 4 healthy chicks were hatched, and the surrogate F_0_ males (n=2) were reared to sexual maturity. Each F_0_ cockerel was then placed in a breeding pen with wild-type (WT) female laying hens and F_1_ eggs were collected daily for analysis.

**Figure 1.**
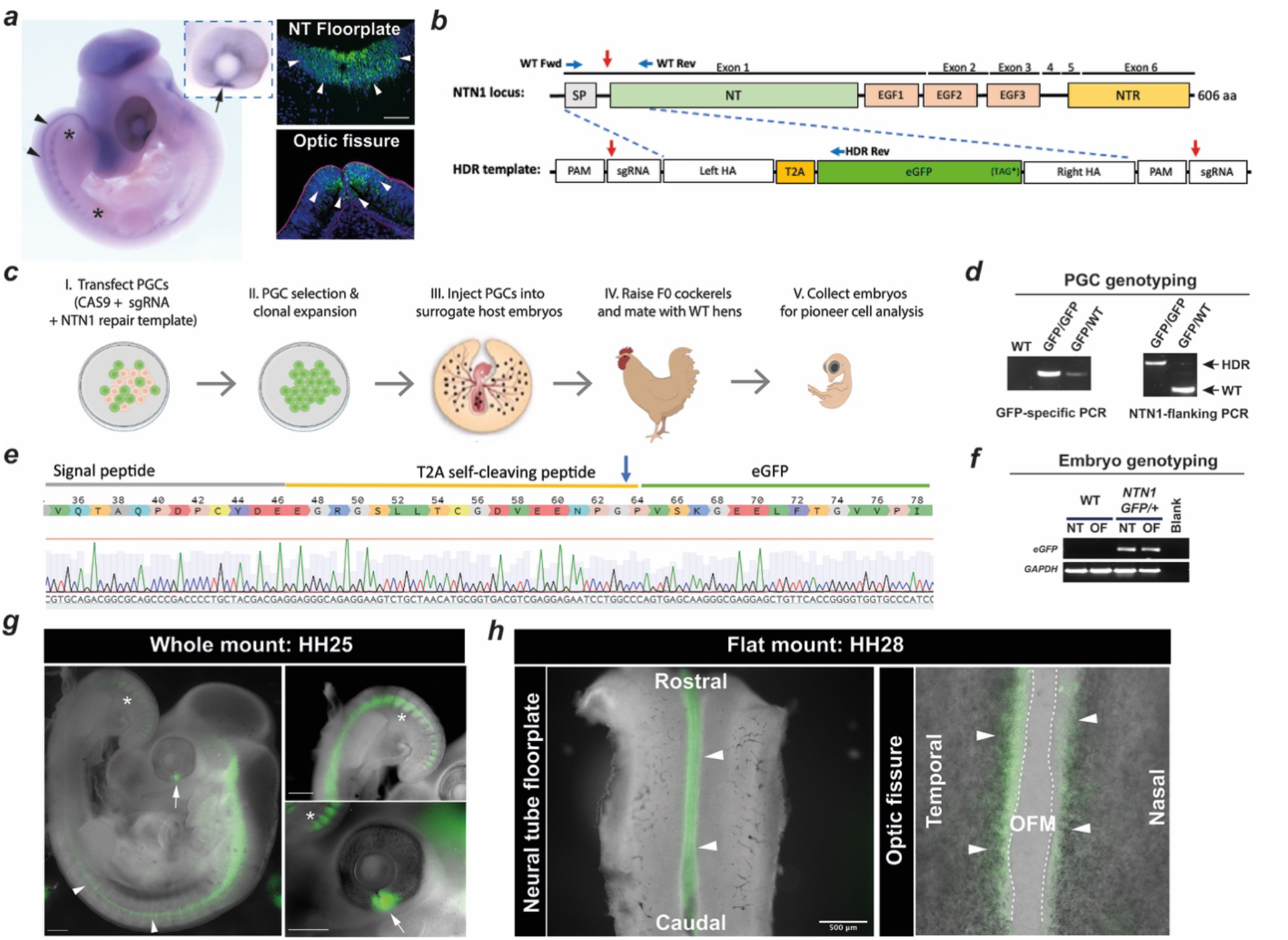
Generation and validation of the *NTN1-T2A-GFP* chicken line. (***a***) Expression of endogenous *NTN1* in wild type chicken embryo in somite regions (between asterisks), neural tube (arrowheads), and in pioneer cell regions at the fissure edges (arrow, inset), and fluorescent *in situ* hybridisation for *NTN1* mRNA (*green*, arrowheads. Scale bar, 50 μm. Blue = DAPI; Magenta = Laminin. (***b***) Schema of genome editing strategy with sgRNA location (red arrow) shown in genome and vector. Abbreviations: EGF, Epidermal growth factor domain; NTR, N-terminal region; PAM, protospacer motif. Blue arrows indicate locations of diagnostic PCR primers. (***c***) Strategy for generating germ-line gene edited *NTN1-T2A-eGFP* chicken line via transfection of PGCs and injection of PGC clones into surrogate host embryos. (***d***) Diagnostic PCRs indicate monoallelic (*NTN1*^*GFP*/WT^) or biallelic (*NTN1*^*GFP*/GFP^) HDR-mediated integration of RTs in clonal PGCs (higher molecular weight band). (***e***) DNA sequence trace and corresponding amino-acid translation showing successful in-frame HDR in clonally propagated PGCs. Arrow indicates predicted T2A self-cleavage site. (***f***) RT-PCR for transgene expression in neural tube (NT) and optic fissure (OF) dissected from WT or *NTN1*^*NTN1-T2A-GFP/+*^ embryos. (***g***) Whole *NTN1*^*NTN1-T2A-GFP/+*^ embryo with GFP expression detected in neural tube (arrowheads), optic fissure (arrows), and somite regions (asterisks). (***h***) Flat-mount *NTN1*^*NTN1-T2A-GFP/+*^ HH28/E6 dissected neural tube (*Left*) opened by cutting dorsally with GFP expression (arrowheads) observed in the floor-plate region, and ventral retina (*Right)* with GFP detected at the edges of the fissure margin (arrowheads).

### *NTN1*^*T2A-GFP*^ replicates endogenous developmental *NTN1* expression and is a pioneer cell reporter

We clearly detected expression of the integrated *NTN1-T2A-GFP* allele in dissected neural tube and optic fissure tissues by RT-PCR of F_1_ embryos (**Figure 1f**). Fluorescence stereomicroscopy revealed GFP expression in *NTN1*^*T2A-eGFP+*^ embryos in the optic fissure, floor-plate of the neural tube, and somite regions, at multiple stages (**Figure 1g-h** and **Supplemental Figure S1**), faithfully recapitulating endogenous *NTN1* expression (**Figure 1a**). We also detected GFP and concomitantly reduced endogenous NTN1 at the protein level in *NTN1*^*T2A-eGFP/+*^ embryo tissues by western blot (**Supplemental Figure S1**), consistent with loss of effective Netrin-1 translation from the gene edited allele. We then tested for embryo viability by macroscopic screening for embryonic defects. the first >150 F1 embryos analysed at embryonic day 9 (∼HH34) or later we observed 99.5 % viability (heartbeat at time of cull, no gross anatomical abnormalities; **Table 1**). All embryos were heterozygous for the NTN1-T2A-eGFP edit (e.g. 100% were *NTN1*^*T2A-eGFP/+*^), consistent with complete colonisation and germ-line activity of the introduced homozygous PGCs in the surrogate F_0_ male gonads (Ballantyne et al., 2021). At day6/HH28, immediately prior to fusion and when the optic fissure margins meet, *NTN1*^*T2A-eGFP* +^ embryos were phenotypically indistinguishable from WT controls and displayed normal anatomy in the optic fissure and no broad ocular abnormalities (**Supplemental Figure S1**). Thus, the NTN1-T2A-GFP line was sufficiently validated to faithfully recapitulate endogenous *NTN1* expression and produce GFP positive (GFP+ve) cells in the optic fissure pioneer cell domain. Furthermore, the introduced *T2A-eGFP* allele caused no observable deleterious phenotypes in heterozygous embryos and in >120 hatched chicks. In combination, these approaches confirmed the reporter lines utility as a system for the study of *NTN1*-expressing cells during normal embryonic development.

**Table 1.**
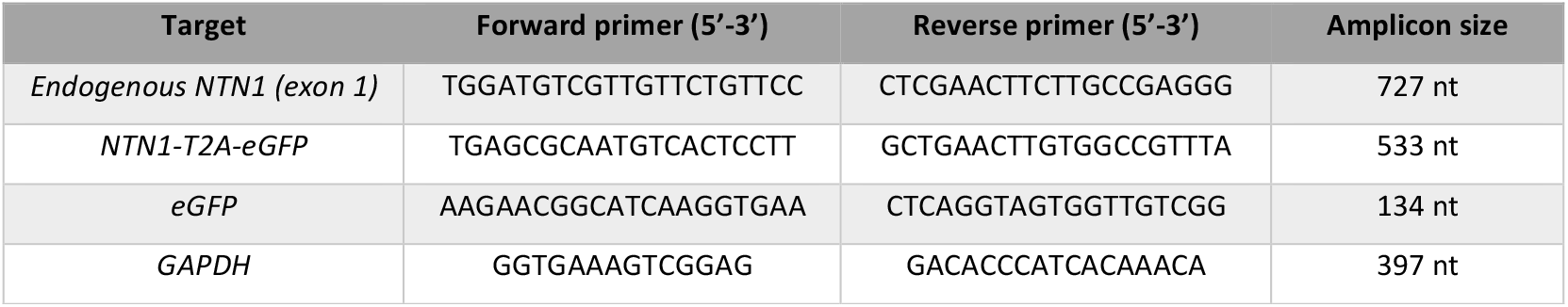
Oligonucleotide primers for allele-specific PCR and RT-PCR.

**Table 2.** Gross phenotyping and viability assessment for *NTN1*^*T2A-eGFP/+*^ heterozygous chicken embryos.

|  |  |
| --- | --- |
| **Fertility** (embryonic development observed) | > **95.5%** (data from 186 eggs) |
| **Viability** (embryos without deformities) | < E9: **97 %** (163/168)<br>< E14: **100 %** (39/39)<br>Hatched chicks: > **95 %** ($n=120$) |

### Isolation of pioneer cells from *NTN1*^*eGFP/+*^ embryonic eyes

The ocular GFP expression was confirmed to the pioneer cell region of the optic fissure (**Figure 1g, Supplemental Figure S1**). We then aimed to isolate these pioneer cells for molecular profiling using a dissection-dissociation and fluorescence activated sorting (FACS) approach (**Figure 2a**). Several protocols exist for generating single cell dissociations from chicken embryonic tissue but were optimised for young chicken embryos, and we found they were not suitable for dissociating ventral chick eye tissue at HH.28 (data not shown). Furthermore, RPE cells can auto-fluoresce at wavelengths that overlap with GFP spectra (Bermond et al., 2020), potentially confounding specific GFP+ve cell isolation by FACS. Thus, to utilise the *NTN1*^*GFP/+*^ embryos for accurate transcriptomic profiling of pioneer cells we first optimised the cell dissociation of dissected fissure margins, and then applied a spectral-sorting FACS strategy to accurately isolate GFP+ve cells (**Figure 2b-c, Supplemental Figure S2**). For cell dissociation, we adapted a method published for mouse eye (Alsing et al., 2022) using a pre-treatment of dissected fissures with hyaluronidase, followed by two rounds of incubation in trypsin/EDTA. Cell suspensions were then analysed by FACS, first defined by the exclusion of debris and dead cells using forward and side scatter, then single cells were selected using side scatter profiling (SSC-A versus SSC-H). We first used wild-type (WT) OFM cell suspensions (i.e. no GFP) to set a conservative fluorescent gating for background fluorescence (**Figure 2b**) and then repeated the process, this time with pooled dissected-dissociated OFMs from *NTN1*^*T2A-eGFP/*+^ embryos (*n* >10 OFMs from independent embryos, per pool). *NTN1*^*T2A-eGFP/*+^ samples were further analysed by spectral flow cytometry to subtract autofluorescence from the populations and stringently define the GFP+ve population (**Figure 2c, Supplemental Figure S2**). We collected cells from both the GFP-ve and GFP+ve fractions from *NTN1*^*T2A-eGFP/+*^ embryos using this method. This approach successfully and reproducibly generated 5 replicate (*n*=5) sample groups each of GFP+ve and GFP-ve cells from the optic fissures of *NTN1*^*T2A-eGFP/+*^ embryos, with each group containing >10,000 cells (**Supplemental Table S1**).

**Figure 2.**
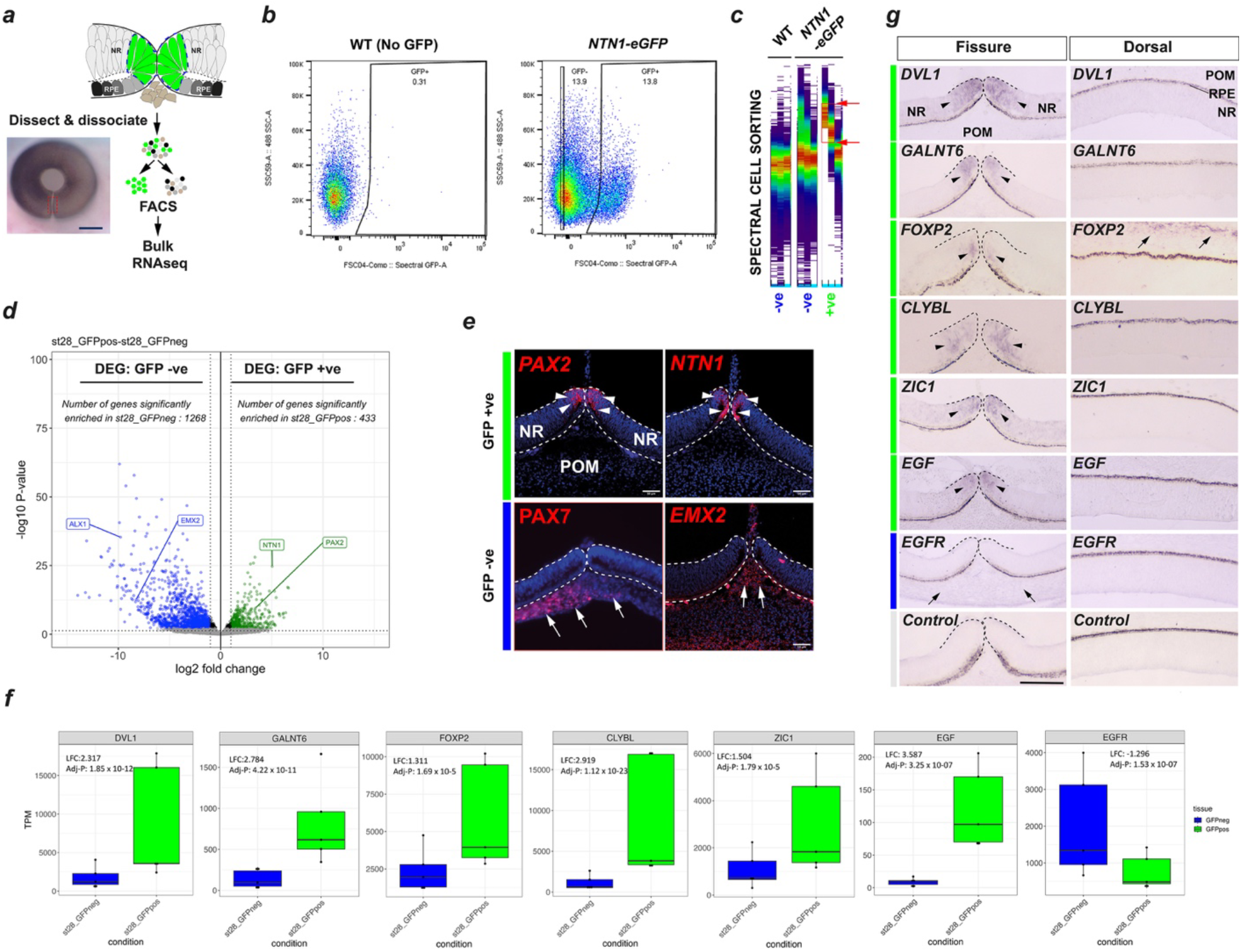
Isolation and transcriptomic profiling of pioneer cells from *NTN1*^*NTN1-T2A-GFP/+*^ reveals novel pioneer cell markers with regulated expression during OFC. (***a***) Schema for optic fissure transcriptomic profiling strategy: manually dissected fissures (red hatched lines) from HH28 *NTN1*^*NTN1-T2A-GFP/+*^ embryos were dissociated and FACS sorted prior to bulk RNA sequencing. (***b***) The GFP+ve and GFP-ve sorted cell populations from *NTN1*^*NTN1-T2A-GFP/+*^ eyes were defined by gating based on using non-fluorescent WT control samples (i.e. no GFP). (***c***) GFP+ve and GFP-ve sorted *NTN1*^*eGFP*^ cells were further separated using spectral flow cytometry to subtract auto fluorescent cells from the GFP+ve population (red arrows). (***d***) Volcano plot of all DEGs defined by -log10 *p*-value and Log2 fold change. There were 433 GFP+ve enriched DEGs (LFC_2_ >1.0) and 1268 DEGs enriched in GFP-ve (LFC_2_ < -0.5). GFP+ve DEGs included known pioneer cell markers *NTN1* and *PAX2*, whereas GFP-ve DEGs included periocular mesenchyme markers *ALX1* and *EMX2*. (***e***) Fluorescence *in situ* hybridisation confirmed *NTN1* and *PAX2* mRNA expression was localised to pioneer cells (arrowheads) at HH28. In contrast *ALX1* and *EMX2* mRNA expression were identified in periocular mesenchyme (arrows). (***f***) Box plots for novel pioneer cell markers identified in this study, with log2 fold change and adjusted-*P* values indicated. (***g***) Colorimetric *in situ* hybridisation for novel OFM genes at HH28 showed pioneer cell specific expression (arrowheads) in the open fissure at HH28 before fusion. Non pioneer cell is expression indicated by arrows.

### Transcriptomic profiling of pioneer cells

We isolated total RNA from the sample groups and generated cDNA libraries for paired-end RNAseq which gave >40 M reads per sample (**Supplemental Table S1**). We aligned reads passing QC to the chicken genome (ENSEMBL, Red Jungle Fowl GRCg6a). PCA analysis of the RNAseq data separated the samples by GFP+ve or GFP-ve for PC1 (62% of variance), and by library preparation for PC2 (29%); i.e. >90% of the variance across the samples could be attributed to either “cell type” or library preparation (**Supplemental Figure S2**). We then performed differential gene expression (DEG) analysis, comparing the two groups (GFP+ve vs GFP-ve) (**Figure 2d**). We found a total of 1701 DEGs (adj *P* < 0.05) across the dataset (**Supplemental Table S2**), with 433 DEGs (LFC ≥ 1) in the GFP+ve cells and 1268 in the GFP-ve cells (LFC ≤1) (**Figure 2d**). To assess broad-scale differences between GFP+ve and GFP-ve, we then assessed enrichment for cell-type specific markers based on gene ontologies (**Supplemental Figure S2**). This analysis showed the GFP-ve cell populations had significantly higher expression of genes in ontology groups for neural crest migration (GO:0001755), epithelial to mesenchymal transition (GO:0001837), mesenchymal development (GO:0060485), neuroepithelial cell differentiation (GO:0060563), and erythrocyte development (GO:0048821), reflecting the enrichment for cells of the periocular mesenchyme, neural retina, and developing hyaloid blood vessels in the dissected tissue and the proportion of these cells compared to pioneer cells in the total dissected material. In contrast, both GFP+ve and GFP-ve showed similar gene expression for the ontology group retinal pigment epithelium development (GO:0003406), alluding to similar transcriptional identities of pioneer cells to RPE and the broadly equivalent cell numbers of each fraction. Furthermore, the GFP+ve enriched gene set included *NTN1* (LFC > 5.01; *p*adj < 6.3E-23) and *PAX2* (LFC > 3.4; adj *P* < 1.1E-07)(**Figure 2d**), both of which had previously been associated with optic fissure closure and the only genes until now shown to have specific expression in pioneer cells (Hardy et al., 2019; Richardson et al., 2019; Trejo-Reveles et al., 2023). In contrast, the periocular mesenchyme marker *EMX2* (LFC < -8.3, padj < 9.9E-11) (Trejo-Reveles et al., 2023) and multiple well-characterised neural crest markers, e.g. *ALX1, PITX1, PITX2, LMX1B, TWIST1, SNAI1*, and multiple members of the *TFAP2* gene family, were all found enriched in the GFP-ve cells (**Figure 2d; Supplemental Table S2**). To confirm these differential gene expression profiles qualitatively, we used fluorescence *in situ* hybridisation on OFM cryosections (***Figure 2e***). This analysis confirmed that both *NTN1* and *PAX2* expression was specific to pioneer cells, while *ALX1* and *EMX2* were both expressed in the periocular mesenchyme, i.e. non-pioneer cells. Thus, these analyses confirmed that the GFP+ve cell population isolated from chicken *NTN1*^*T2A-eGFP/+*^ ventral eye tissue reflects the first specific isolation of optic fissure pioneer cells from the surrounding ventral eye environment and provided the foundation for further molecular profiling of pioneer cells during fusion.

### *NTN1*^*T2A-eGFP/*+^ pioneer cell profiling reveals multiple novel OFC candidates

We next selected pioneer-specific DEGs for qualitative analysis by *in situ* hybridisation in wild-type chicken embryonic eye tissue at fusion stages, based on their TPM enrichment values (**Figure 2f**): *DVL1*, encoding Dishevelled-like 1 (LFC = 2.317, adj *P* = 1.85 E-12), *GALNT6* encoding polypeptide N-acetylgalactosaminyltransferase 6 (LFC = 2.784, adj *P* = 4.22 E-11), *FOXP2* encoding the forkhead box protein P2 transcription factor (LFC = 1.311, adj *P* = 1.69 E-05), *CLYBL* encoding citramalyl-CoA lyase (LFC2.919, adj *P* = 1.12 E-23), the Zic-family transcription factor *ZIC1* (LFC = 1.504, adj *P* = 2.01 E-05), and the epidermal growth factor encoding gene *EGF* (LFC = 3.59, adj *P* = 3.25 E-07). To the best of our knowledge neither of these genes had previously been identified as expressed in the optic fissure or associated with ocular coloboma or embryonic tissue fusion, thus represented interesting novel OFC/coloboma candidate genes. Our analysis showed that all were specifically expressed in the pioneer cell domain of the optic fissure and were not detected in the broader ventral or dorsal retina, although *FOXP2* was detected in the dorsal periocular mesenchyme (**Figure 2g**). *FOXP2* also showed the most restricted expression domain within this panel, with signal in a region of the OFM located in the small group of cells adjacent to the folding point where pioneer cells are apically-tethered prior to fusion (Chan et al., 2021; Hardy and Rainger, 2023). Of additional interest was that the EGF receptor encoding gene *EGFR* was enriched in the GFP-ve DEG data (LFC = -1.29; Adj *P* = 1.92E-08) and our mRNA *in situ* indicated this expression was localised to the periocular mesenchyme and was therefore expressed in a distinct tissue region to the site of expression of its ligand *EGF*, which was pioneer cell enriched (**Figure 2g**). To assess these genes during active fusion we performed additional *in situ* hybridisation on serial or semi-serial sections at HH30 and found that *GALNT6, CLYBL, DVL1* and *ZIC1* all showed expression in the pioneer cells prior to and during fusion, with immediate downregulation of expression in the nascently fused seam (**Supplemental Figure S2**). Thus, using this approach we validated the use of *NTN1*^*T2A-eGFP/+*^ as a tool to identify genes with enriched expression in the pioneer cell region, and based on the restricted expression profiles for those we tested, we propose these represent multiple new candidates for functional roles in optic fissure closure.

### Pathway enrichment analyses indicate novel signalling pathways in pioneer cells

To identify active molecular pathways in pioneer cells which may drive fusion, we took all 433 NTN1-GFP+ve DEGS and performed gene ontology (GO) analyses using online *in silico* analysis tools (**Figure 3a**). This approach revealed enrichment for a range of processes and biological pathways of relevance to OFC and epithelial fusion, the most significant (adj *p* <0.05) of which included: *plasma membrane, signalling by receptor tyrosine kinases, signal transduction, signalling or receptor-ligand binding by G-coupled protein receptors, PI3K-AKT signalling, and various extracellular, plasma membrane, and ligand-receptor processes* (**Figure 3a, Supplemental Table S2**). We then plotted the expression values for all GFP+ve DEG genes associated with the KEGG pathway term PI3K/AKT and found 17 genes in this dataset, including the tyrosine kinase receptor *MET* and its ligand *HGF* (**Supplemental Figure 3**). We also performed STRING enrichment analysis to highlight protein-protein interactions within the GFP+ve DEG set. This revealed multiple similar networks to the ontology analyses but highlighted that there were significant interactions from a range of molecules that all had MET as their core interaction partner (**Figure 3b; Supplemental Figure S3**). In combination, these *in silico* analyses suggested that HGF/MET and PI3K/AKT related signalling pathways may be active in chicken pioneer cells during fusion. HGF and MET are the secreted ligand and canonical receptor, respectively, for hepatocyte growth factor signalling (HGF, “scatter factor”), which upon ligand-receptor interactions induces a range of cellular responses, including proliferation, cell survival, and invasive growth (Birchmeier and Gherardi, 1998; Trusolino and Comoglio, 2002). Confirming our *in silico* analyses, the RNAseq data showed that *MET* expression was significantly enriched in pioneer cells (LFC = 3.53; adj *P* = 2.7E-14; **Figure 3c**). Although *HGF* expression in the GFP+ve DEGs did not reach our significance thresholds (*HGF* LFC = 0.94; adj *P* = 0.43), absolute TPM levels of *HGF* revealed that its expression was higher in GFP+ve compared to GFP-ve cells (**Figure 3c**). We then performed dual fluorescence *in situ* hybridisation for *MET* and *HGF* and observed that both genes were indeed specifically expressed in the OFM, but their expression was localised to distinct domains along the proximal-distal axis in the pre-fusing fissure at both HH25 and HH28 (**Supplemental Figure S3** and **Figure 3d**). At HH28, immediately prior to fusion, *MET* expression was clearly identified in RPE and neural retinal cells at the pioneer cell region in the medial OFM where fusion will subsequently initiate but was absent from the distal (iris) region where fusion does not occur (**Figure 3d** and **Supplemental Figure S3**). In contrast, at the same stage *HGF* expression was detected broadly in the ventral neural retina in the distal OFM (not RPE) but was not detected in the medial pioneer cell region (**Figure 3d**). Thus, despite both genes being expressed in the optic fissure margins, at no point along the proximal distal axis did *HGF* and *MET* expression overlap in the pre-fusing and HH28 OFM. Fusion is active and dynamic at HH30 in the chicken OFM, with all phases of fusion present: open fissure, fusion plate and a fused seam (Hardy et al., 2019). Fluorescence *in situ* hybridisation at this stage revealed that expression of *MET* was still evident within the pioneer cells at the apposed fissure margins, in the fusion plate, and in the immediately adjacent (within 100 μm of the fusion plate) fused seam (**Figure 3e**). In contrast, HGF expression was not detected in the in the pioneer cells but was expressed in a small domain at the junction between pioneer cells and the RPE (**Figure 3f**). Finally, to provide direct evidence for activity of the HGF/MET pathway in the fusing OFM, we used immunofluorescence against phosphorylated MET protein (phos-Tyr1234/phos-Tyr1235) in the HH30 chicken fissure (**Figure 3g**). In accordance with the adjacent expression of *MET* and *HGF* at this stage leading to autophosphorylation of the receptor at cell surfaces of ligand expressing cells (Trusolino et al., 2010), we observed fluorescence signal in the apical region of pioneer cells immediately prior to fusion and in the fusion plate but were unable to detect signal in the fusion seam >100 μm from the fusion plate. In combination, our data provide plausible evidence for fusion-specific activity of the HGF/MET pathway in pioneer cells.

**Figure 3.**
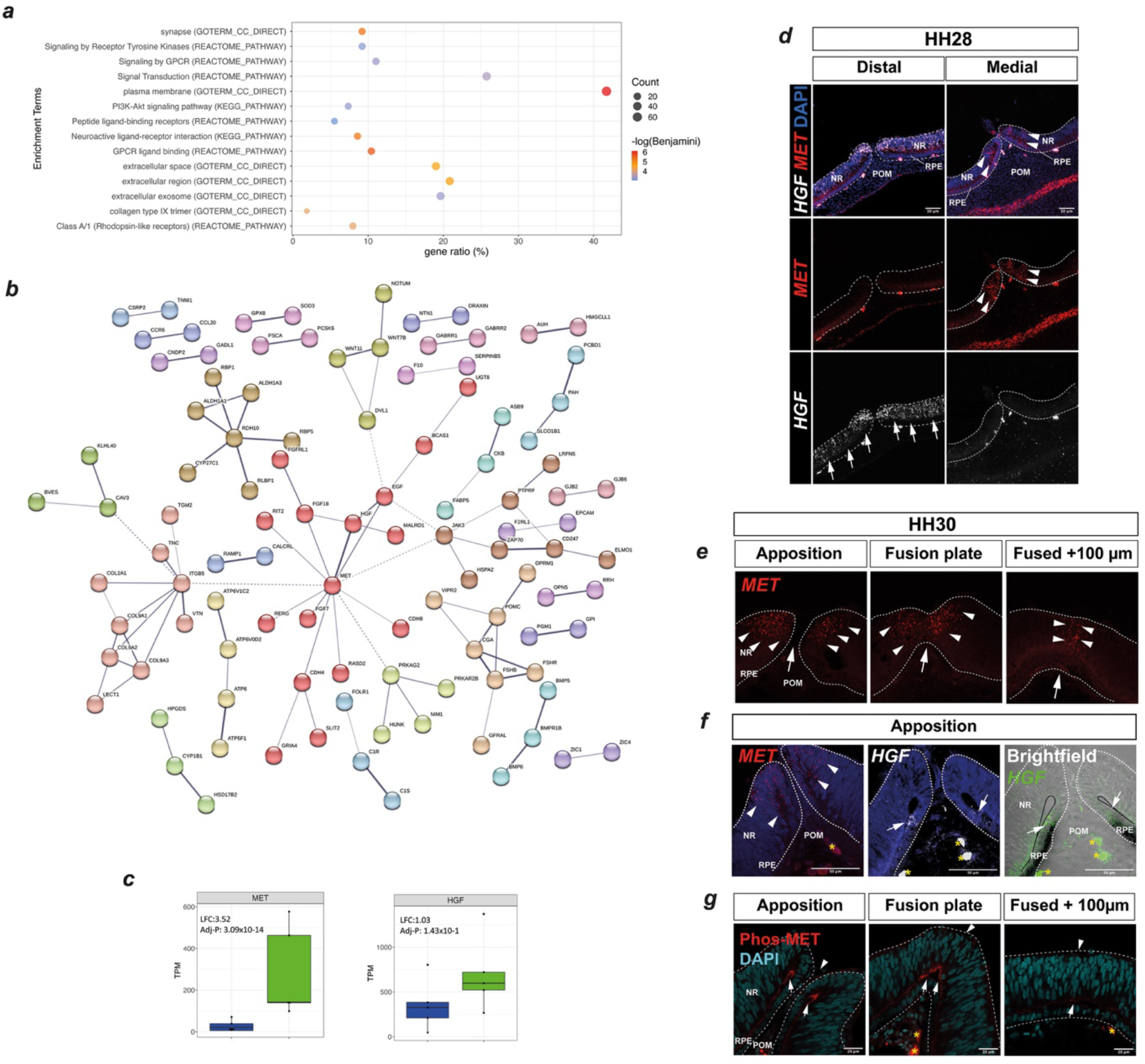
Pathway analyses profiling reveals HGF/Scatter factor activity during active fusion. (***a***) Consolidated plots showing gene ratios and counts for significant ontology enrichment terms (adj *p* <0.05) associated with GFP+ve enriched DEGs using KEGG Pathway, Reactome Pathway, and GO TERMs. (***b***) STRING Network analysis of protein-protein interactions within the GFP+ve DEG set indicated MET at the centre of interactions and directly connected to its ligand HGF. (***c***) Box and whisker plots of absolute gene expression level (transcripts per million, TPM y-axis) for *MET* and *HGF* in GFP+ve (green plots) and GFP-ve cells (blue plots). (***d***) Fluorescence *in situ* hybridisation analyses at prefusion stage HH28 for *MET* (filled arrows) and *HGF* (yellow open arrows) in the distal and medial optic fissure margins. Note non-overlapping expression domains. (***e***) Fluorescence *in situ* hybridisation analyses during active fusion (HH30) indicated specific expression of *MET* (arrowheads) in the neural retina pioneer cell domain. (***f***) Comparison of *MET* and HGF expression in the fusing OFM revealed *HGF* expression (arrows) in cells at the junction of the RPE and pioneer cells, which was immediately adjacent to the *MET* expressing pioneer cells. (***g***) Immunofluorescence analysis for phosphorylated MET receptor protein (Phospho-Tyr1234/Tyr1235) identified activity at the folding point of the OFM connecting the RPE border cells to the apical regions of pioneer cells (arrows). Signal was not markedly reduced in the fused seam at 100 μm from the fusion plate in the same eye. Large arrowheads in ***e*** indicate the midline of the OFM; asterisks represent auto fluorescent red blood cells. Abbreviations: RPE, Retinal pigmented epithelium; POM, periocular mesenchyme; NR, neural retina.

### Broader utility of the *NTN1-T2A-eGFP* chicken line

Given there are multiple other areas of *NTN1* expression in the developing embryo (Hardy et al., 2019; Nishitani et al., 2017; Salminen et al., 2000b), *NTN1*^*T2A-eGFP/+*^ embryos could be similarly useful to profile other developing tissues with *NTN1* expression, such as semi-circular epithelial canals during inner ear formation, the floor plate of the neural tube (e.g. rostral versus caudal NT transcriptional profiles), and hinge regions of the developing pharyngeal arches. In addition, recent evidence suggests *NTN1* is expressed in corneal epithelial cells in response to scratch wounding (Han et al., 2020), during vascularisation and kidney development (Honeycutt et al., 2023), and in other contexts including a range of metastatic tumours (Chaturvedi and Murray, 2021; Cirulli and Yebra, 2007). Therefore, we propose that a similar dissociation-FACS approach could be used to specifically collect *NTN1* expressing cells for a range of multi-omics techniques to enable profiling and expedite understanding of their biology. Thus, this new stable *NTN1-GFP* chicken line is an excellent tool for *NTN1* biology and optic fissure closure research.

## METHODS

### Generation of NTN1-T2A-GFP PGCs

CRISPR guides were designed using Chop Chop (https://chopchop.cbu.uib.no/), selecting the optimal guide sequence with the lowest predicted off-target cut sites in exon 1 of *NTN1* (NM_205419.1). Male primordial germ cells (PGCs) were isolated from day 2.5 Hy-Line chick embryonic blood samples and selectively grown using previously published protocols (Whyte et al., 2015). PGCs were transfected using Lipofectamine 2000 (Thermo) according to the manufacturer protocol with the CRISPR/Cas9 plasmid pX458 (Ran et al., 2013) containing the *NTN1*-specific sgRNA sequence: 5’-GACCCCTGCTACGACGAGCA-3’. A second plasmid was co-transfected that contained an HDR repair template flanked at the 5’- and -3’ ends by the same *NTN1* sgRNA recognition and sequences for a T2A self-cleaving peptide in-frame with enhanced GFP protein coding sequence, flanked on both sides with 100 bp homology arms specific to either side of the *NTN1* targeted region locus in exon 1. The full DNA sequence of the insert is available in **Supplemental S1**. Transfected PGCs were treated with puromycin and then FACS sorted to individual cells and seeded into 96-well plates following the protocol of Ballantyne et al (Ballantyne et al., 2021). Of 120 single cell cultures, we eliminated dead clones as well as the fastest and slowest growing clones. For the remaining clones, we cryopreserved cell stocks for injections and screened the clonal populations by genomic DNA extraction and sequence specific PCR, using oligonucleotide primers specific for the endogenous *NTN1* locus and sequences specific for the HDR repair template (**Table 1**). HDR clones were confirmed by Sanger sequencing to identify in-frame HDR events with no indels at the recombined locus. Confirmed homozygous NTN1-T2A-eGFP PGC clonal populations were propagated and then frozen for subsequent embryo injections.

### Generation and phenotypic analysis of NTN1-T2A-eGFP chicken line

All animal management, maintenance and embryo manipulations were carried out under UK Home Office licence PP0018931 at the University of Edinburgh. Experimental protocols and studies were approved by the Roslin Institute Animal Welfare and Ethical Review Board Committee. A monoclonal male targeted PGC line was thawed and injected into 12 iCaspase9 surrogate host embryos according to previously published protocols (Ballantyne et al., 2021). Briefly, to remove endogenous germ cells in the host chicken embryos 1.0 μl of 25 mM B/B (in DMSO) (Takara Bio) was added to 50 μL PGC suspensions before injection. Chicks (*n*=4/12) were hatched, and the two males (2/4) were raised to sexual maturity. Each male was placed in breeding pens with 6-7 WT Hy-Line female layers and eggs collected on a weekly basis. Eggs were incubated at 37.5°C up to 14 days and assessed for viability (normal embryonic development with no obvious phenotypic anomalies) and fertility (evidence of embryonic development). Observations were taken at 6, 9, or 14 days of incubation. Eye measurements were performed using brightfield images and a measurement graticule. Eye diameters of stage-matched WT and *NTN1*^*-T2A-eGFP/+*^ embryos were measured and compared at day 9. The horizontal and vertical (coronally) dimensions of the globe were measured using digital callipers (resolution; 0.01 mm) and these measurements were averaged per eye and compared between genotypes. Graphpad Prism was used to perform one-way ANNOVA statistical tests. Tissues were collected from all embryos, and genomic DNA was extracted using PureLink Genomic DNA Mini Kit (Thermo Scientific) according to the manufacturer’s instructions. PCR and Sanger sequencing was performed using *NTN1* and *GFP* specific primers to confirm inheritance of the integrated alleles. For non-quantitative RT-PCR optic fissure and neural tube tissues were dissected and total RNA was isolated using Trizol reagent (Ambion Life Sciences) according to manufacturer instructions. Total RNA (1 μg) was reverse transcribed using oligo-dT primers with Superscript III (Thermo) according to the manufacturer’s instructions. PCR was then performed using *eGFP* or *GAPDH* sequence specific primers and amplicons were run on 1 % agarose gels. Primers are listed in Table 1.

### *In situ* gene expression and immunofluorescence analyses

WT embryos were collected and staged (Hamburger and Hamilton, 1992) and fixed for 2 hours in 4% Paraformaldehyde (PFA) in phosphate buffered saline (PBS), pH 7.0, then rinsed in PBS. For whole mount in situ hybridisation and cryosection RNAscope, embryos were processed according to Hardy et al., 2019 and Trejo-Reveles et al., 2023 (*NTN1, EMX2, PAX2*) following the RNAscope Multiplex Fluorescent v2 Kit (ACD, Biotechne) protocol for frozen tissue. New RNAscope probes were designed for *MET* (20ZZ probe named Gg-MET targeting 218-1255 of NM_205212.3; Cat No. 1248471-C1) and *HGF* (20ZZ probe named HGF/SF targeting 260-1365 of NM<u>_</u>001030370.5; Cat No. 1273221-C3). For fluorescent *in vivo* analysis of GFP expression in *NTN1-T2A-eGFP* embryos we imaged freshly dissected embryos immersed in PBS using a Zeiss StereoLumar V12 Fluorescence Stereomicroscope. For immunofluorescence analysis we used 20 μm cryosections incubated with antibodies against GFP (#ab5450; Abcam; 1:500 dilution), Laminin (#3H11; DSHB; 1:5), or Anti-Phos (Tyr1234/Tyr1235) MET (CST #3077; 1:60), overnight at 4°C in 0.1 % BSA-PBS with 0.1 % Tween. Alexa-Fluor fluorescently conjugated secondary antibodies (Thermo) were used at 1:1000 dilutions. F-actin (Phalloidin-488 nm; Thermo; 1 in 200) and DAPI (4′,6-diamidino-2-phenylindole; Thermo; used at 1:1000) were also used as fluorescent counter stains and mounted with coverslips using Fluorsave (Merck). Imaging was performed on Zeiss LSM-800 confocal microscopy and images managed using Zen Black software (Zeiss) and FIJI (Schindelin et al., 2012).

### Western Blotting and reverse-transcriptase PCR

Whole optic fissures or neural tube tissues were manually dissected from HH28 *NTN1*^*T2A*-eGFP*/+*^ embryos into lysis buffer (Yung et al., 2015). Protein levels were estimated using BCA assays against known standards and mixed with SDS-lamelli loading buffer and heated to 95°C for 10 mins, then cooled immediately on ice. Proteins (20 μg) were loaded and run on 4%-12% Bis-acrylamide SDS gels and transferred onto nitrocellulose membranes before blocking in 5% milk TBS-Tween (0.01%) solution for 1 hour at room temperature. Membranes were then incubated in primary antibodies overnight at 4°C with gentle agitation, then rinsed in TBS-Tween and incubated in HRP-conjugated secondary antibodies for 1h at room temperature. Membranes were then washed in TBS-Tween before detection with Pierce ECL Western Blotting Substrate (Thermo).

### Cell dissociation and isolation

Embryos were removed from their eggs and staged according to Hamburger and Hamilton (Hamburger and Hamilton, 1992). Heads were removed and transferred to a new dish in HBSS (Gibco) on ice. Optic fissure tissue was dissected, ensuring removal of cornea, lens and sclera, and as much excess tissue from the nasal and temporal non-OFM retina as possible. Pooled OFMs were placed in Lo-Bind 1.5 mL tubes (Eppendorf) and rinsed twice in fresh HBSS, using a benchtop centrifuge to collect sample tissue at the base of tubes by brief (15 second) spins each time. All HBSS was removed then replaced with 900 μL fresh HBSS and 100 μL hyaluronidase added (10 mg/ml; Sigma) and incubated at 37°C for 10 minutes. Samples were then placed on ice and OFM tissue allowed to settle before removal of the hyaluronidase and replaced with 900 μL fresh cold HBSS. Samples were then centrifuged with a mini microfuge for 10 seconds and left on ice for a further two minutes, then briefly centrifuged for 10 seconds and all HBSS removed. 500 μL of Trypsin/EDTA 0.25% (Gibco) was added and samples were incubated for 5 minutes at 37°C. During this time, samples were triturated by pipetting up and down 15 times every 2 min using P200 low bind tips. Dissociation was assessed by taking 10 μL of sample combined with trypan blue and loaded onto a Neubauer cell counting chamber. 500 μL of 20 % FBS-HBSS (Sigma) was added to samples to stop the trypsin reaction and tubes were centrifuged at 500 x*g* for 2 minutes at room temperature supernatant removed. Samples were then resuspended in 500 μL 0.05 % trypsin for maximum 4 minutes at 37°C, pipetting up and down several times using P200 low bind tips every 2 minutes. 500 μL HBSS was added, and samples were centrifuged at 500 x*g* for 2 minutes at room temperature and supernatant was removed. The final dissociated cell preparations were then resuspended in 500 μL HBSS, with 10 μL removed for cell counting and viability assessment.

### Fluorescence cell sorting and spectral unmixing

Dissociated cells were transferred to a Flow Cytometry Tube (BD) and filtered through the cap to remove highly aggregated cells. An extra 500 μL HBSS was added to rinse the filter and maximize the cell collection. Dissociated cells were separated into GFP positive (GFP+ve) and GFP negative (GFP-ve) populations using a Bigfoot cell sorter and Sasquatch (SQ) software version 1.19.40295.3 (Thermo). The sorting strategy involved excluding debris, clumps and dead cells using FSC-A and SSC-A. Single cells were selected from cell aggregates by SSC-A vs SSC-H, followed by FSC-A vs FSC-H. The GFP+ve gate was defined based on the absence of cells in scatter plots of dissociated cells taken from WT chicken optic fissure margin dissections with no fluorescence. This was always performed before FACS with NTN1^T2A-eGFP^ containing samples. To sort the NTN1^T2A-eGFP^ dissociated cells we used 10,000 cells to generate the spectral signature that enabled autofluorescence subtraction from the background population to specifically define the GFP+ve population (**Supplemental Figure S2**). Lo-Bind 1.5 mL tubes (Eppendorf) collection tubes were pre-coated in PBS with of 5 % Bovine Serum Albumin (Sigma Aldrich) and sorted cells were collected in 800 μl of the same solution prior to being centrifuged at 500 x*g* for 5 minutes and supernatants removed prior to RNA extraction.

### RNAseq transcriptomic profiling

Dissociated GFP+ve and GFP-ve cells were immediately processed for total RNA isolation using PicoPure RNA isolation kit (Articus, Thermo), applying the DNaseI step. Total RNA was assessed on the Agilent 2100 Electrophoresis Bioanalyser Instrument (Agilent Technologies Inc) and RNA 6000 Pico chips for quantity, quality, and integrity of total RNA. Libraries were prepared from 7 ng of total-RNA samples using the NEBNext Single Cell/Low Input RNA Library Prep Kit for Illumina (NEB), according to the provided protocol. Libraries were quantified by fluorometry using the Qubit dsDNA HS assay and assessed for quality and fragment size using the Agilent Bioanalyser with the DNA HS Kit. Fragment size and quantity measurements were used to calculate molarity for each library pool. Sequencing was performed on the NextSeq 2000 platform (Illumina Inc) using NextSeq 1000/2000 P2 Reagents (200 Cycles) v3. Libraries were combined in an equimolar pool of 6 based on Qubit and Bioanalyser assay results and run over a P2 flow cell. PhiX Control v3 (Illumina Inc) was spiked into each run at a concentration of 1%. Basecall data was converted into FASTQ files using BaseSpace (Illumina) and downloaded to local drives at the University of Edinburgh. All sequence files are publicly available at NCBI GEO with accession GSE275781.

### Bioinformatic analyses

Sequencing quality was evaluated using FASTQC (version 0.11.4). Sequencing adaptors were trimmed using Cutadapt (Martin, 2011) (version 1.16) and Trimmgalore (version 0.4.1). Trimmed sequences were aligned to the chicken genome (galgal6) using STAR (version 2.7.1a) to generate binary alignment and map (BAM) files, which were then coordinate sorted and indexed using Samtools (Li H, Handsaker B, Wysoker A, Fennell T, Ruan J, Homer N, Marth G, Abecasis G, Durbin R, 2009) (version 1.6). A count matrix of transcripts annotated in Ensembl (version 105) was subsequently generated from BAM files using featureCounts part of the Subread package (Liao, Y., Smyth, G. K., & Shi, 2013) (version 2.0.1). Differential gene expression and principal component analysis was performed using DESeq2 (Love, M. I., Huber, W., & Anders, 2014) in R (version 4.1.0), with batch correction:

~~~
batch<-factor(c(rep(“batch1”,4),rep(“batch2”,(ncol(countdata)-4))))
coldata <-data.frame(row.names=colnames(countdata),
condition,tissue,batch)
dds_all <-DESeqDataSetFromMatrix(countData=count_matrix,
colData=coldata_subset,
design= ∼tissue+batch)
~~~

Adjusted *P*-values were determined from comparisons of differentially expressed genes between GFP-positive and GFP-negative populations using Benjamini and Hochberg correction (Benjamini, Y. and Hochberg, 1995). Genes with a log2 fold change ≥ 1.0 and an adjusted P-value ≤0.05 were considered as upregulated GFP+ve associated differentially expressed genes (DEGs), while those with a log2 fold change of ≤ -1.0 and an adjusted P-value of ≤ 0.05 were considered as downregulated or GFP-ve DEGs in the analysis. Volcano plots of DEGs were drawn using ggplot2 package in R. For generation of TPM quantification files, STAR-aligned BAM files were passed to Salmon (version 1.3.0) for quasi-mapping (Patro et al., 2017). Transcript annotation for Salmon was based on Ensembl (version 105) and was extracted from Ensembl gtf file using gffreads (version 0.12.7). Salmon quantification files were then loaded on R (version 4.1.0) using Tximport (version 1.22.0). Boxplot indicating median TPM value of individual gene were created using an in-house script and ggplot2 (version 3.4.2). For TPM heatmaps, Z-score of genes of interest were calculated individually across different samples using an in-house script and plotted using ComplexHeatmap (version 2.10.0).

### Cell type grouping

Gene names related to the following GO-terms were extracted from AmiGO (version 2.5.17) (Carbon et al., 2009): erythrocyte development (GO-0048821), neuroepithelial cell differentiation (GO-0060563), retinal pigment epithelium development (GO-0003406), neural crest cell migration (GO-0001755), mesenchyme development (GO-0060485), and epithelial to mesenchymal transition (GO-0001837). Ensembl ID (version 105) of the corresponding gene names for each GO term category were identified, and only those genes with valid *Gallus gallus* ensembl IDs were used in the analysis. Mean TPM value of each gene was then calculated in GFP+ve and GFP-ve samples, respectively, and those with mean TPM values < 0.1 in both GFP-ve and GFP+ve samples were filtered out. For each GO term category, mean TPM values of genes within the category were plotted as violin plots using ggplot2 (version 3.4.2). Paired *t*-test was performed for the pairwise comparison of mean TPM for each gene between GFP+ve and GFP-ve samples using `stat_compare_means` function by ggpubr (version 0.6.0).

### Ontology enrichment and network analyses

To identify enrichment of pathways and gene ontology groups we used PANTHER Overrepresentation Test (Released 20221013): GO biological process complete, and DAVID analysis tools: KEGG PATHWAY and GO TERM. Specifically, we used GFP+ve DEGs for PANTHER GO biological process complete with “Client Text Box Input” (*Homo sapiens*) as our Analysed List, and the background “Reference List” *Homo sapiens* (all genes in database). Test Type: Fisher’s Exact. Annotation Version and Release Date: GO Ontology database DOI: 10.5281/zenodo.7942786 Released 2023-05-10. For DAVID, we used the Functional Annotation Tool to analyse KEGG_PATHWAY and GOTERM_BP_DIRECT using only those GFP+ve DEGs with Official Gene IDs and with *Homo sapiens* as a background list and ENSEMBL gene IDs as inputs. Lists of GO terms were ranked and plotted (-log10(pval)) with cut off values p < 0.001) (Huang da W, Sherman BT, 2009; Sherman et al., 2022). Plots were prepared using ggplot2 (version 3.4.2). For STRING (Version 11.5) protein-protein network analysis, we inputted all DEGs in the GFP+ve selection (log2 fold change ≥ 1.0 and an adjusted P-value ≤0.05) as gene names and used the following settings: Full STRING network; hide disconnected nodes; “evidence” to define network edges; “Textmining, Experiments, Databases, Co-expression and Neighbourhood” criteria selected as active interaction sources; Minimum required interaction score - Medium > 0.400. Data shown in **Supplemental Table S2**. Figure prepared with extra node prediction by STRING, MCL clustering with inflation parameter of 1.5, for visualization proteins without linkage were dropped out. A total of 10 nodes were added, leading to the node (i.e. protein) number of 283, number of edges (i.e. functional association linkage) was 96 and a PPI enrichment score of 6.73 × 10^−7^.

## ACKNOWLEDGEMENTS

We wish to acknowledge the support and insightful scientific discussions with Françoise Helmbacher and Flavio Maina (Aix-Marseille Université), Brian Brooks (NIH, National Eye Institute), and Megan Davey and Tom Burdon (Roslin Institute). We are also highly grateful to the staff at the National Avian Research Facility at Roslin Institute for chicken line husbandry, and our colleagues in the Bioimaging facility for help with confocal imaging and cell sorting.

## DATA AVAILABILITY

The authors have deposited all sequencing files on GEO (https://www.ncbi.nlm.nih.gov/geo/) with reference number GSE275781.

