## Supplemental data for "A stable *Netrin-1* fluorescent reporter chicken reveals cell-specific molecular signatures during optic fissure closure"

### Supplemental Table S1. Repair template sequence, FACS cell number yield, RNAseq quality metrics.

### Supplemental Table S2. DEG lists and ontology & enrichment outputs.

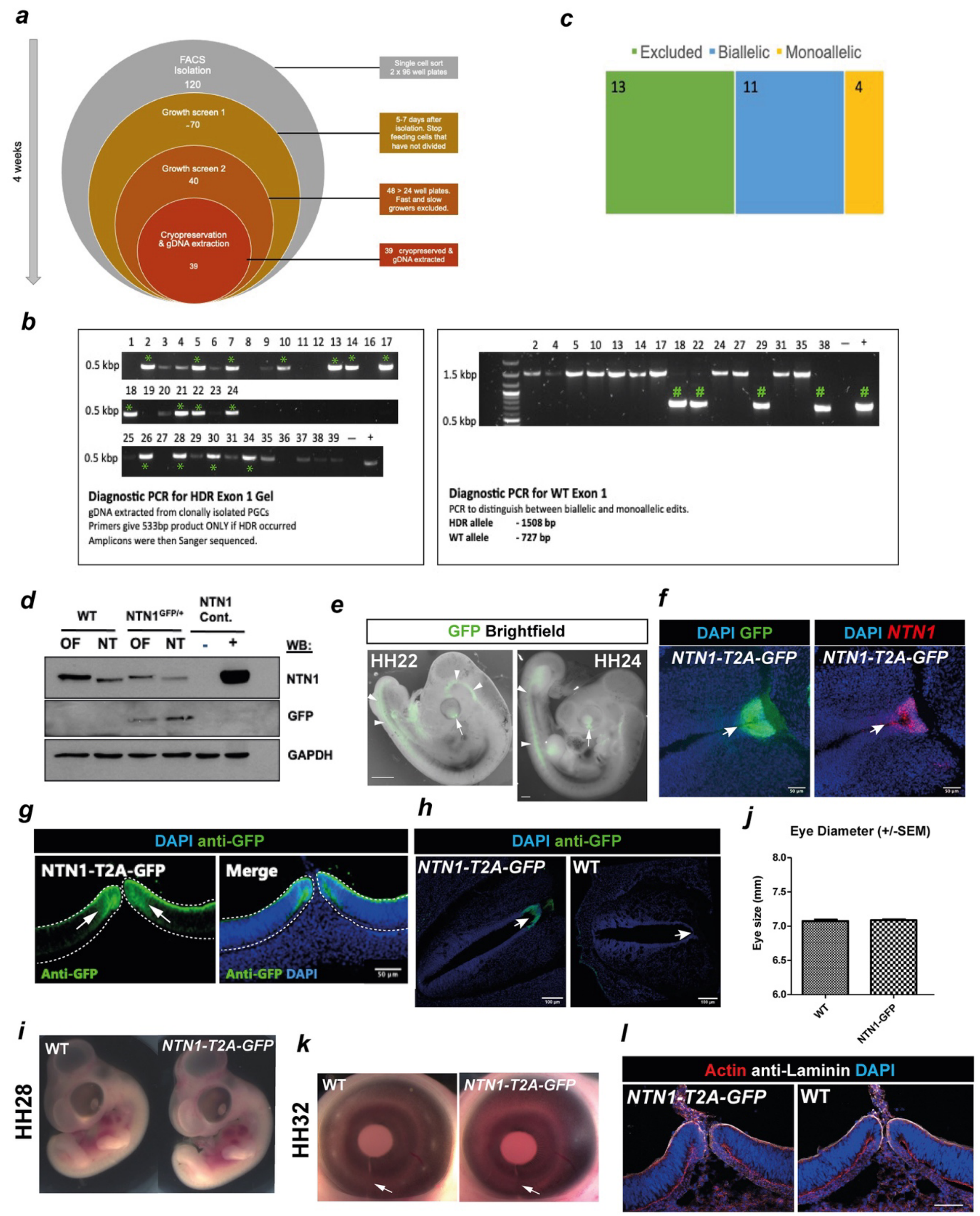

**Supplemental Figure S1.** (***a***) Schema for clonal *NTN1-T2A-GFP* PGC generation and validation. Transfected PGCs were single-cell sorted by FACS into individual wells and grown, then screened twice for successful and normal growth, then frozen with genomic DNA extracted for diagnostic PCR and Sanger sequencing. (***b***) Diagnostic PCRs to select unambiguous amplicons for successful HDR (left) and distinguish biallelic and monoallelic HDR events. (***c***) Summary statistics for cryopreserved PGCs – 11x clonal lines contained in-frame biallelic HDR knock in of T2A-eGFP with no mismatches, and 4x were monoallelic. (***d***) GFP and endogenous NTN1 proteins were detected in $NTN1^{NTN1\text{-}T2A\text{-}GFP/+}$ and WT embryos from NT and OF tissues by western blot. GAPDH was used as a loading control Flp-In T-Rex 293 cells with NTN1 stably integrated were used as a positive control, the parent line was used a negative control. (***e***) Whole mount stereomicroscope images show GFP localisation in the developing embryo at HH22 and HH24. Arrowheads, neural tube; arrows, optic fissure. (***f***) GFP localisation in the floorplate of the $NTN1^{NTN1\text{-}T2A\text{-}GFP/+}$ neural tube at HH28, followed by RNAscope fluorescent *in situ* hybridisation for *NTN1* on the same sample, indicating co-localisation. (***g,h***) GFP detection (arrows) by anti-GFP immunofluorescence in the HH28 optic fissure pioneer cell region (***g***) and floorplate of the neural tube (***h***). (***i***) $NTN1^{NTN1\text{-}T2A\text{-}GFP/+}$ stage-matched embryos were grossly phenotypically normal compared to WT at HH28. (***j***) Eye sizes of $NTN1^{eGFP/+}$ embryos were phenotypically normal compared to WT. (***k***) Fusion of the optic fissure (arrows) was complete by HH32 (~ day 8) in all $NTN1^{NTN1\text{-}T2A\text{-}GFP/+}$ embryos and controls. (***l***) Heterozygous $NTN1^{NTN1\text{-}T2A\text{-}GFP/+}$ embryos displayed normal optic fissure anatomy at HH28 compared to WT, illustrated by distribution of basement membrane (Laminin), cell (F-actin) and nuclei (DAPI).

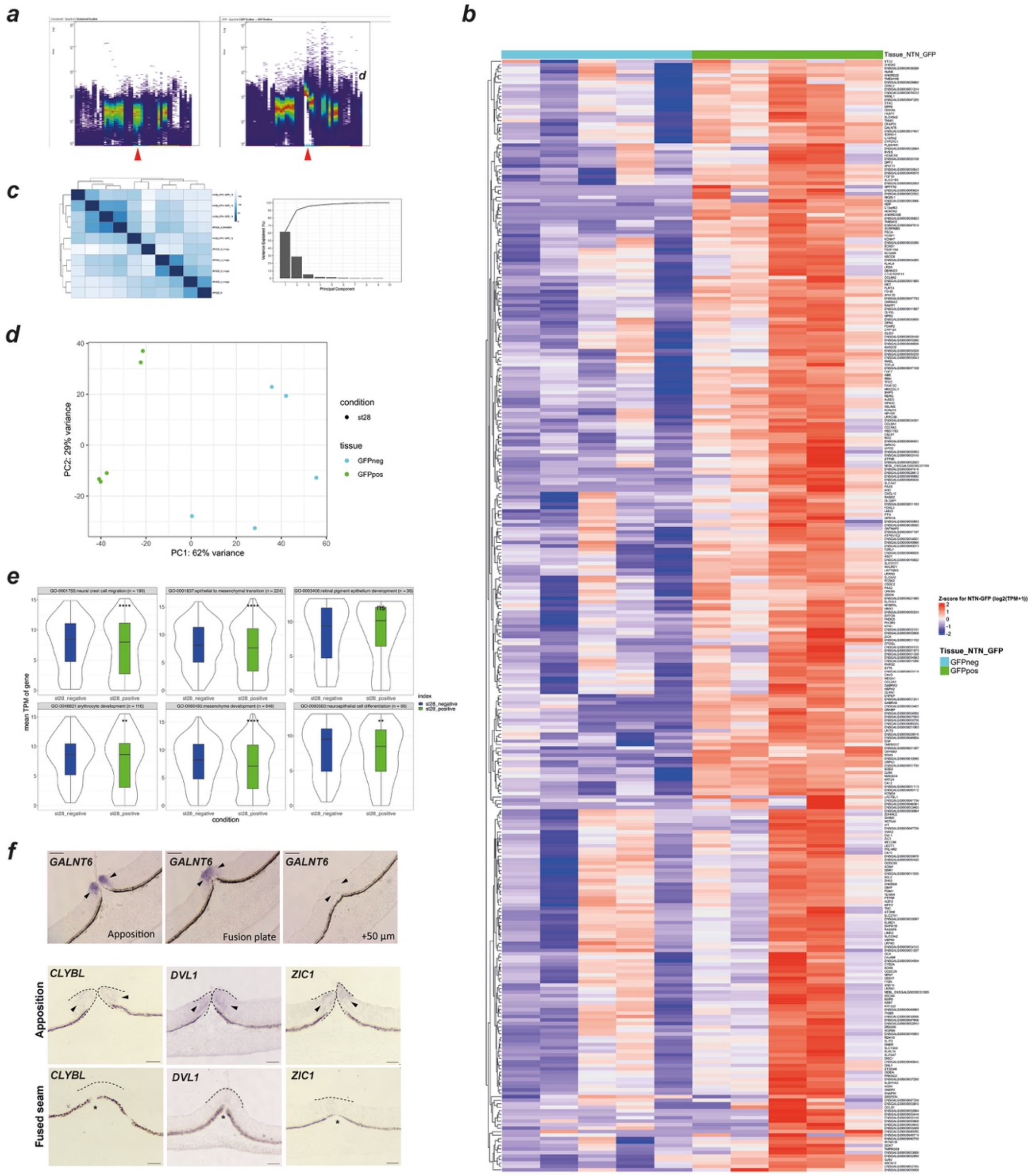


**Supplemental Figure S2.** (***a***) Spectral profiling of WT and *NTN1*$^{NTN1\text{-}T2A\text{-}GFP/+}$ FACS. GFP+ve and GFP-ve sorted *NTN1*$^{eGFP}$ cells were separated using spectral flow cytometry to subtract auto fluorescent cells from the GFP+ve population. Note the clear spectral shift in GFP+ve population compared to GFP-ve population (red arrowheads) in the blue excitation wavelengths. (***b***) Heat map showing expression profiles for the GFP+ve enriched genes across all 10 samples. (***c***) Correlation matrix showing relatedness of all RNAseq samples. Sample ID: JR1222_5_Mar2023 is *NTN1*$^{NTN1\text{-}T2A\text{-}GFP/+}$ whereas JR1222_9 is WT. Scree plot of all principal components. Note PC1 and PC2 account for >90% of the variance between all samples. (***d***) Principal component analysis plot of RNAseq data for GFP+ve and

GFP-ve samples (*n*=5, per group). PC1 separated samples by GFP+ve vs GFP-ve (62% variance). (***e***) Violin plots showing distribution of expression levels (log10 TPM) for genes in ontology groups associated with cell types. Significance values identified using paired *t*-test: ** p< 0.01; ***p < 0.001; **** p <0.0001; ns = non-significant. (***e***) Overview of number of GFP-enriched genes ascribed to KEGG terms associated with MET. (***f***) *in situ* hybridisation for a subset of identified pioneer cell specific genes in the actively fusing HH30 optic fissure shows temporally regulated expression, with specific expression observed (arrowheads) in the pioneer cell domain during apposition, then absence of detectable mRNA in the fully fused seam. Serial sections were used for *GALNT6* analysis and similar temporally regulated fusion-specific expression was observed for *CLYBL*, *DVL1* and *ZIC1*.

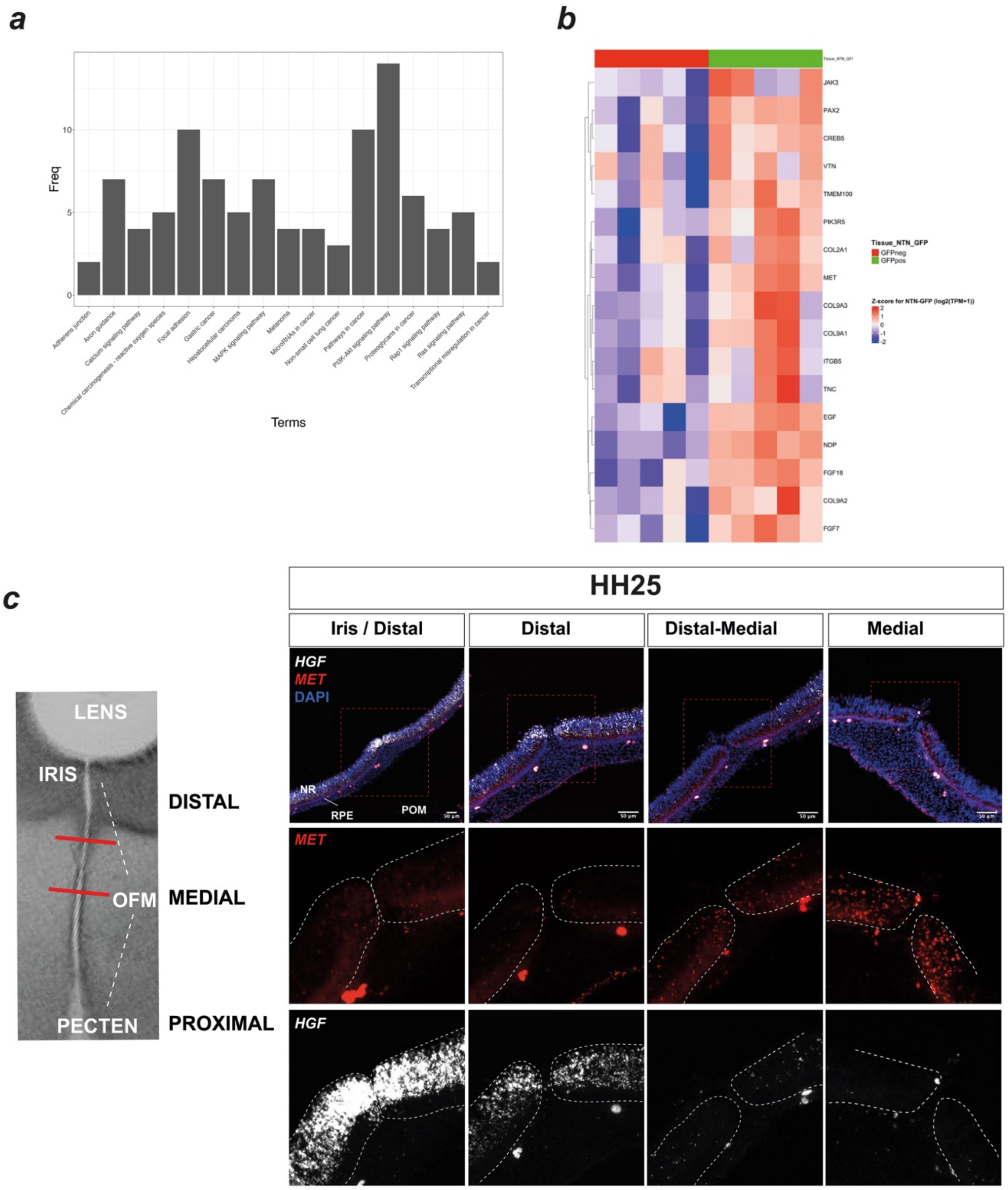


**Supplemental Figure S3.** (***a***) frequency plots for genes listed within the ontology terms associated with the GFP+ve pioneer cell enriched geneset. (***b***) Heatmap showing PI3K/Akt pathway associated genes enriched in GFP+ve pioneer cells. (***c***) Combined fluorescence *in situ* hybridisation for *HGF* and *MET* with semi-serial sections shows non-overlapping domains of expression in the HH25 OFM: *HGF* is expressed in the distal OFM, whereas *MET* is expressed in the medial OFM at the future point of fusion.
